# Functional characterization of RebL1 highlights the evolutionary conservation of oncogenic activities of the RBBP4/7 orthologue in *Tetrahymena thermophila*

**DOI:** 10.1101/2020.11.07.372946

**Authors:** Syed Nabeel-Shah, Jyoti Garg, Alejandro Saettone, Kanwal Ashraf, Hyunmin Lee, Suzanne Wahab, Nujhat Ahmed, Jacob Fine, Joanna Derynck, Marcelo Ponce, Shuye Pu, Edyta Marcon, Zhaolei Zhang, Jack F Greenblatt, Ronald E Pearlman, Jean-Philippe Lambert, Jeffrey Fillingham

**Affiliations:** Department of Chemistry and Biology, Ryerson University, 350 Victoria St., Toronto M5B 2K3, Canada; Department of Biology, York University, 4700 Keele St., Toronto, M3J 1P3, Canada; Department of Computer Sciences, University of Toronto, Toronto, M5S 1A8, Canada; Donnelly Centre, University of Toronto, Toronto, M5S 3E1, Canada; Department of Molecular Genetics, University of Toronto, Toronto, M5S 1A8, Canada; SciNet HPC Consortium, University of Toronto, 661 University Avenue, Suite 1140, Toronto, ON M5G 1M1, Canada; Department of Molecular Medicine, Cancer Research Center, Big Data Research Center, Université Laval, Quebec, Canada; CHU de Québec Research Center, CHUL, 2705 Laurier Boulevard, Quebec, G1V 4G2, Canada

**Keywords:** RBBP4/7, Epigenetics, Chromatin, Cancer analysis, Gene expression, Functional proteomics, Histones, MuvB/DREAM complex, Ciliates

## Abstract

Retinoblastoma-binding proteins 4 and 7 (RBBP4 and RBBP7) are two highly homologous human histone chaperones. They function in epigenetic regulation as subunits of multiple chromatin-related complexes and have been implicated in numerous cancers. Due to their overlapping functions, our understanding of RBBP4 and 7, particularly outside of Opisthokonts, has remained limited. Here, we report that in the ciliate protozoan *Tetrahymena thermophila* a single orthologue of human RBBP4 and 7 proteins, RebL1, physically interacts with histone H4 and functions in multiple epigenetic regulatory pathways. Functional proteomics identified conserved functional links associated with *Tetrahymena* RebL1 protein as well as human RBBP4 and 7. We found that putative subunits of multiple chromatin-related complexes including CAF1, Hat1, Rpd3, and MuvB, co-purified with RebL1 during *Tetrahymena* growth and conjugation. Iterative proteomics analyses revealed that the cell cycle regulatory MuvB-complex in *Tetrahymena* is composed of at least five subunits including evolutionarily conserved Lin54, Lin9 and RebL1 proteins. Genome-wide analyses indicated that RebL1 and Lin54 (Anqa1) bind within genic and intergenic regions. Moreover, Anqa1 targets primarily promoter regions suggesting a role for *Tetrahymena* MuvB in transcription regulation. RebL1 depletion decreased cellular viability and altered the expression of selected targets. Consistent with observations in glioblastoma tumors, RebL1 depletion suppressed DNA repair protein Rad51 in *Tetrahymena*, thus underscoring the evolutionarily conserved functions of RBBP4/7 proteins. Our results suggest the essentiality of RebL1 functions in multiple epigenetic regulatory complexes in which it impacts transcription regulation and cellular viability.

## Introduction

Eukaryotic chromatin provides a means to compact the genome as well as contributing to regulating DNA-mediated processes, such as replication, recombination, repair, and transcription (1). Histone chaperones and chromatin remodelers, as well as histone posttranslational modifications (PTMs), play pivotal roles in chromatin-related processes (2). For example, multiple histone chaperones, such as nuclear autoantigenic sperm protein (NASP) and anti-silencing factor 1 (Asf1), have been reported to function in the transport pathway of newly synthesized H3/H4 (3, 4). Furthermore, chromatin assembly factor-1 (CAF1) and histone-regulator-A (HIRA) function in DNA replication-dependent (RD) and replication-independent (RI) chromatin assembly processes to deposit either H3–H4 or H3.3–H4, respectively (5, 6). ATP-dependent chromatin remodelers, on the other hand, mobilize DNA around nucleosomes and function in transcription regulation (7). For example, the ‘Nucleosome Remodeling and Deacetylase’ (NuRD) complex that is widely present at gene promoter regions and enhancer elements plays an important role in regulating transcription and genome integrity, as well as cell cycle progression (8, 9). Finally, histone PTMs are important for chromatin assembly, as well as gene expression regulation (10). Newly synthesized H4 histones are di-acetylated at lysine (K) 5 and K12 residues, which has been shown to be important for chromatin assembly (11). These deposition-related H4K5/12ac marks are installed by the HAT1-complex, which is composed of at least two subunits, including catalytic Hat1 and histone-binding WD40 repeat protein Hat2 (RBBP4 in humans) (12, 13). In *Saccharomyces cerevisiae* an additional nuclear subunit called Hif1 (human NASP) also interacts with the HAT1-complex at roughly stoichiometric levels (14). Whereas the deposition-related H4 PTMs are conserved across species (15), the corresponding enzymatic complex has not been well described in evolutionarily distant protozoan lineages.

Retinoblastoma-binding proteins 4 and 7 (RBBP4 and RBBP7) (Cac3 and Hat2 in budding yeast, respectively), also known as Retinoblastoma Associated protein 48 (RbAp48) and 46 (RbAp46), respectively, are two highly homologous histone chaperones (~90% sequence identity) with similar structures (16) that were initially identified as retinoblastoma-associated proteins (17, 18). Both proteins contain WD40-repeats forming a seven-bladed β-propeller fold, which is thought to function as a protein interaction scaffold in multiprotein complexes (16, 19). While human and mouse encode both homologues, only a single homologue of RBBP4 and RBBP7 has been identified in some organisms, including *Caenorhabditis elegans* (Lin53) and *Drosophila melanogaster* (p55) (20–22). RBBP4 and RBBP7 histone chaperones are components of several multi-protein complexes that are involved in diverse chromatin-related functions, including chromatin assembly and remodeling, histone post-translational modifications and gene expression regulation (23, reviewed by 24). For example, RBBP4 and RBBP7 have been identified as subunits of the NuRD complex (25), switch independent 3A (Sin3A) complex (26), and polycomb repressive complex 2 (PRC2) (26–28). Furthermore, RBBP4 (Cac3) has been found to be a subunit of the CAF1 complex (29, 30), and RBBP7 (Hat2) has been reported to be an essential component of the HAT1-complex (31). Within multi-subunit protein complexes, RBBP4 and RBBP7 are thought to function as chromatin adaptors via their direct interaction with H3/H4 (16, 32).

RBBP4 also functions as a component of transcriptional regulator ‘multi-vulva class B’ (MuvB)-complex (33–35). The mammalian core MuvB complex, composed of five subunits including Lin9, Lin37, Lin52, Lin54, and RBBP4, is considered to be the master regulator of cell-cycle-dependent gene expression (33). The composition of the core MuvB remains unaltered during different cell cycle stages, whereas its interaction partners change, which reverses its function from a transcriptional repressor to an activator (33, 36, 37). The core MuvB interacts with E2F4/5, DP1/2 and p130/p107 proteins to form the repressor ‘dimerization partner (DP), RB-like, E2F and MuvB’ (DREAM) complex, which represses the G1/S and G2/M genes during G0 and early G1 (35, 36). The DREAM-specific proteins dissociate during late G1, and the MuvB core then forms an interaction with the B-MYB transcription factor during S phase (35, 38). The interaction of B-MYB with MuvB is necessary for FOXM1 recruitment during G2 (38). These B-MYB-MuvB (MMB) and FOXM1-MuvB complexes stimulate the expression of genes that peak during G2 and M phases (34). Among the MuvB core subunits, Lin54 is a DNA-binding protein and has two tandem cysteine-rich (CXC) domains that share sequence similarity with Tesmin, a testis-specific metallothionein-like protein (39, 40). Lin54 directly binds to DNA by recognizing the ‘cell cycle genes homology region’ (CHR) element (41–43). The CHR is considered to be the central promoter element required for the cycle-dependent regulation of G2/M phase-expressed genes (43). Several studies have shown that the DREAM-complex localizes to its target promoters via E2F/DP and LIN-54 DNA sequence motifs (44–46). In addition to Lin54, DREAM also likely comes in contact with nucleosomes by utilizing RBBP4 via its interaction with histones (reviewed by 33). Furthermore, *Drosophila* p55 (RBBP4) has been shown to be important for DREAM-mediated gene repression (47).

It is increasingly recognized that epigenetic aberrations contribute to tumor initiation and development (48). As such, it is not surprising that RBBP4 and RBBP7 have been extensively implicated in many cancers and are now being pursued as valuable drug targets (reviewd by 24). For example, RBBP7 is upregulated in many cancers, including non-small cell lung cancer, renal cell carcinoma, and breast cancer (49–51). Similarly, RBBP4 has been found to be overexpressed in hepatocellular carcinoma and acute myeloid leukemia (52, 53). In human glioblastoma (GBM) tumor cells, knockdown of RBBP4 causes sensitization of tumor cells to temozolomide (TMZ) chemotherapy and supresses the expression of several DNA repair genes, including RAD51, a key enzyme of the homologous recombination repair pathway (54).

Despite their critical role in multiple epigenetic pathways, the function of RBBP4 and RBBP7 and their associated protein complexes has not been well documented outside of the Opisthokonta. *Tetrahymena thermophila*, a unicellular ciliate protozoan, is an excellent experimental system to study evolutionarily conserved chromatin-related processes, including gene expression and cell cycle regulation (55–58). *Tetrahymena* has two structurally and functionally distinct nuclei, a germ-line diploid micronucleus (MIC) and a polyploid somatic macronucleus (MAC), that are physically separated within the same cell (59). The MAC essentially regulates all the gene expression during vegetative growth, whereas the MIC ensures stable genetic inheritance (60). Both the MAC and MIC arise from the same zygotic nucleus during the sexual reproduction (conjugation) of the cell (60). During conjugation, two developing nuclei undergo substantial chromatin alterations, including DNA rearrangements and removal of ‘internally eliminated sequences’ (IES), resulting in functionally and structurally different MAC and MIC with distinct epigenetic states (61–63). Many of the epigenetic regulatory pathways in *Tetrahymena* that ensure the establishment of distinct chromatin states within the two nuclei are widely conserved across eukaryotes (64).

Here, we utilized functional proteomics and genomics approaches to characterize the single orthologue of human RBBP4 and RBBP7, RebL1, in *Tetrahymena*. Iterative proteomic analyses uncovered the composition of multiple *Tetrahymena* epigenetic regulatory complexes, such as the CAF1-, Sin3A/Rpd3-and MuvB-complexes, which co-purified with RebL1 during vegetative growth and conjugation. Furthermore, ChIP-seq analysis of RebL1 and Lin54 (Anqa1), combined with previously reported RNA-seq data (65), implicated *Tetrahymena* MuvB in transcription regulation. Consistently, the knockdown (KD) of *REBL1* significantly decreased cellular viability and altered the expression profile of some functionally critical genes, including *RAD51*. Overall, our results revealed a general role of RebL1 in multiple transcriptional regulatory pathways important for cell cycle and chromatin regulation in *Tetrahymena*.

## Results

### RebL1 physically interacts with histone H4

We sought to identify the *Tetrahymena* RBBP4 and RBBP7 histone chaperones by first identifying the interaction partners for the core histone H4, one of their predicted partners. The *Tetrahymena* genome contains two core histone H4s encoded by *HHF1* and *HHF2* (66). We generated *Tetrahymena* cell lines stably expressing *HHF1* and *HHF2* with a C-terminal FZZ epitope tag from their endogenous MAC loci. The FZZ epitope tag comprises 3×FLAG and two protein A moieties separated by a TEV cleavage site, which can be utilized in affinity purification experiments and indirect immunofluorescence (IF), as well as genome-wide studies. The successful expression of the endogenously tagged proteins was confirmed by Western blotting analysis in whole cell extracts (WCEs) prepared either from the *HHF1*-and *HHF2*-FZZ expressing cells or untagged wild-type *Tetrahymena* (Supplemental figure S1A). Similar to the core histone H3 (67), our IF analysis indicated that during vegetative growth HHF1- and HHF2-FZZ localize to both MAC and MIC (Supplemental figure S1B). This observation suggests that the FZZ-tagged histones remain functionally competent, as previously reported (58, 68). Since *HHF1* and *HHF2* encode identical proteins, we used only the *HHF1*-FZZ cells (designated as H4) for our downstream analyses.

Using our established pipeline (69, 70), we performed affinity purifications (AP) coupled to tandem mass spectrometry (AP-MS) in biological replicates to identify the interaction partners of H4-FZZ. To eliminate any possible indirect associations mediated by DNA or RNA, we treated the cell extracts with a promiscuous nuclease (Benzonase) prior to the affinity pull-downs to remove RNA and DNA from our samples. The MS data were scored using the SAINTexpress algorithm, which utilizes semiquantitative spectral counts for assigning a confidence value to individual protein-protein interactions (PPIs) (71). SAINTexpress analysis of the H4-FZZ AP-MS data filtered against several control purifications identified 18 high-confidence co-purifying proteins (≤ 1% false discovery rate [FDR]) (Figure 1A; Supplemental Table S1). The H4-FZZ interaction partners included histone chaperones Asf1 and Nrp1, two heat shock Hsp70s, an Importinβ6, MCM-complex proteins including Mcm-2 and Mcm-6, FACT-complex subunit Spt16 (Cet1), Poly [ADP-ribose] polymerase 6 (PARP6) and Hat1, all of which have been shown to function in histone regulation and/or co-purify with H3 (58, 69, 72, 73).

**Figure 1:**
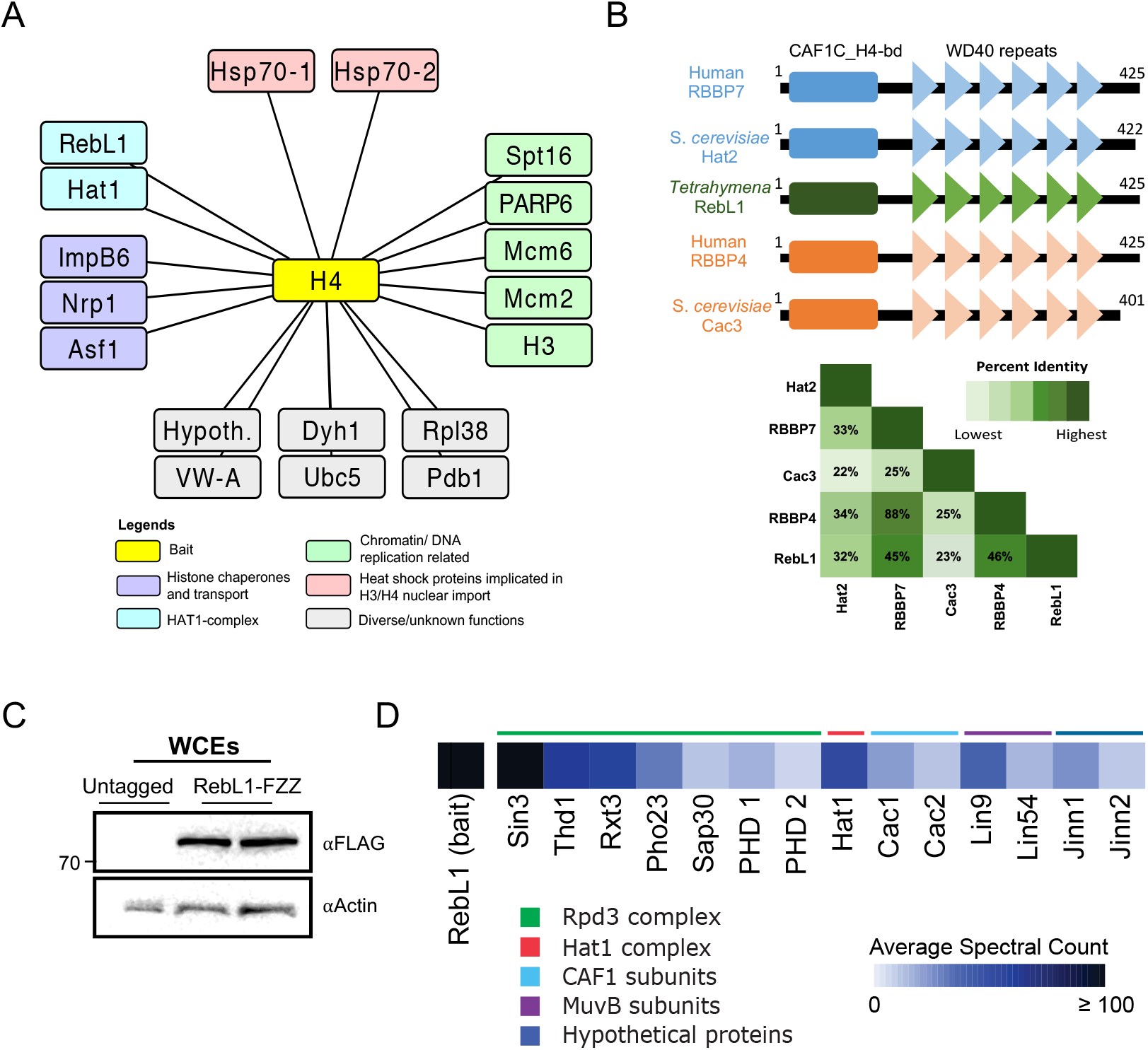
Identification and proteomic characterization of RebL1 in *Tetrahymena*. **A:** Network representation of high confidence (FDR≤0.01) H4 co-purifying proteins. Note: Hypothetical (Hypoth.) is used to indicate proteins that lack annotation on *Tetrahymena* genome database. See Supplemental Table S1 for complete AP-MS results. **B:** Comparative domain analysis of *Tetrahymena* RebL1 protein against human and budding yeast orthologues. Overall sequence identity among the orthologues is shown as half-square rectangle (bottom). **C:** Western blotting analysis using whole cell lysates prepared from two different mating type *Tetrahymena* cells expressing RebL1-FZZ. The blot was probed with the indicated antibodies. **D:** Heatmap diagram of high confidence (FDR≤0.01) RebL1 co-purifying proteins during vegetative growth of *Tetrahymena*. Definition of the color code is indicated. See Supplemental Table S2 for complete AP-MS results.

Finally, we identified TTHERM_00688660 as one of the high-confidence (FDR ≤ 1%) H4 co-purifying proteins which shared sequence similarity with human RBBP4 and RBBP7. The co-purification with H4 of an RBBP4/7-like protein (RebL1, ***Re***tinoblastoma ***B***inding protein 4/7-***L***ike 1) is consistent with previous reports describing human and budding yeast orthologs to function as histone chaperones (74). We subsequently investigated TTHERM_00688660 as a putative RBBP4/7 *Tetrahymena* orthologue in greater detail.

### RBBP4/7 is conserved in *Tetrahymena*

WD40-repeat family proteins are encoded by multiple genes in most organisms (19, 75). We examined the *Tetrahymena* genome for the existence of additional RBBP4/7-like proteins with predicted WD40-repeats. Using the human RBBP4 and RBBP7 proteins to search against the *Tetrahymena* genome, we identified two proteins, including TTHERM_000709589 and the H4 co-purifying TTHERM_00688660 (RebL1). Both identified proteins have conserved domain architecture with an N-terminal H4-binding domain followed by six WD40 repeats (Figure 1B; Supplemental figure S1C). Multiple sequence alignment analysis, however, indicated that TTHERM_000709589, which we named as Wdc1 (***<u>WD</u>***40 domain ***<u>c</u>***ontaining 1), shares only weak sequence identity (~18%) with its human and yeast counterparts; in comparison, RebL1 had an overall ~46% sequence identity (Figure 1B). It is worth noting that *Tetrahymena* also encodes another WD40 repeat protein, TTHERM_00467910, which appears to be an orthologue of *S. cerevisiae* RRB1 (known as GRWD1 in humans) (Supplemental figure S1C), as assessed by reciprocal BLAST searches. RRB1 is known to function in ribosome biogenesis (76), making TTHERM_00467910 an unlikely candidate to be a RBBP4/7 orthologue.

To further categorize the identified WD40 proteins in *Tetrahymena*, we carried out a phylogenetic analysis and observed that RebL1 and Wdc1 are related to RBBP4/7-family of histone chaperones (Supplemental figure S1D). Wdc1 and *Arabidopsis thaliana* Msi4 clustered together, distinctly within the RBBP4/7 group on the phylogenetic tree, suggesting that these proteins might be functionally divergent (Supplemental figure S1D). In contrast, TTHERM_00467910 was found within well differentiated clade representing the RRB1 protein family (Supplemental figure S1D). Collectively, we conclude that RebL1 and Wdc1 are related to the RBBP4/7-family of histone chaperones in *Tetrahymena*, whereas TTHERM_00467910 belong to a phylogenetically distinct WD40 sub-family.

### RebL1 interacts with diverse chromatin-associated protein complexes

To test the hypothesis that TTHERM_00688660 is a *bona fide* RBBP4/7 orthologue, we generated endogenously tagged *Tetrahymena* cell lines of two different mating types stably expressing *REBL1* with a C-terminal FZZ epitope tag from its native MAC locus (Figure 1C). Next, we performed AP-MS analysis in vegetative *Tetrahymena* cells using RebL1-FZZ as a bait. Application of SAINTexpress identified at least 14 high-confidence (FDR≤0.01) interaction partners of RebL1 during vegetative growth (Figure 1D; Supplemental Table S2). These interaction partners include the putative orthologue of *S. cerevisiae* Hat1, as well as TTHERM_00309890 and TTHERM_00219420, which shared sequence similarity with the human p150 and p60 (*S. cerevisiae* Cac1 and Cac2) subunits of the CAF1 complex, respectively (Figure 1D). These observations highlight the potentially conserved heterotrimeric and heterodimeric subunit composition of the CAF1- and HAT1-complexes, respectively. Next, we identified seven RebL1 co-purifying proteins, including Sin3, Thd1, Rxt3, Pho23, Sap30 and two PHD-containing proteins that, based on similarity with the putative *S. cerevisiae* orthologues, appear to define a *Tetrahymena* Rpd3 histone deacetylase (HDAC) complex (Figure 1D). The high-confidence RebL1 interactions also included putative orthologs of the Lin54 (TTHERM_00766460) and Lin9 (TTHERM_00591560) subunits of the mammalian MuvB-complex (Figure 1D). An additional two unannotated (or hypothetical) proteins, TTHERM_00227070 and TTHERM_00147500 (named as Jinn1 and Jinn2, respectively), co-purified with RebL1 and lacked any recognizable domains and/or sequence similarity to any known protein.

As RebL1 expression is highest after meiosis, a state that is marked by a series of rapid post-zygotic nuclear divisions (Supplemental figures S1E and S2), we investigated how its interactome was remodeled during sexual development of *Tetrahymena*. To do so, we performed AP-MS in conjugating cells harvested 5 h post-mixing. After scoring the individual PPIs using SAINTexpress, we detected a total of 30 reproducible, high-confidence interaction partners of RebL1 during conjugation (Supplemental Table S3). All the RebL1 interaction partners identified during vegetative growth were also detected as high-confidence hits in the conjugating *Tetrahymena* (Figure 2A). Functional enrichment analysis of the 30 RebL1 interaction partners identified gene ontology (GO) terms enriched for chromatin organization, transcription regulation and histone modifications (Figure 2B). Among the RebL1 interaction partners exclusively identified during conjugation were four putative MYB-like transcription factors (TFs) (Figure 2C).

**Figure 2:**
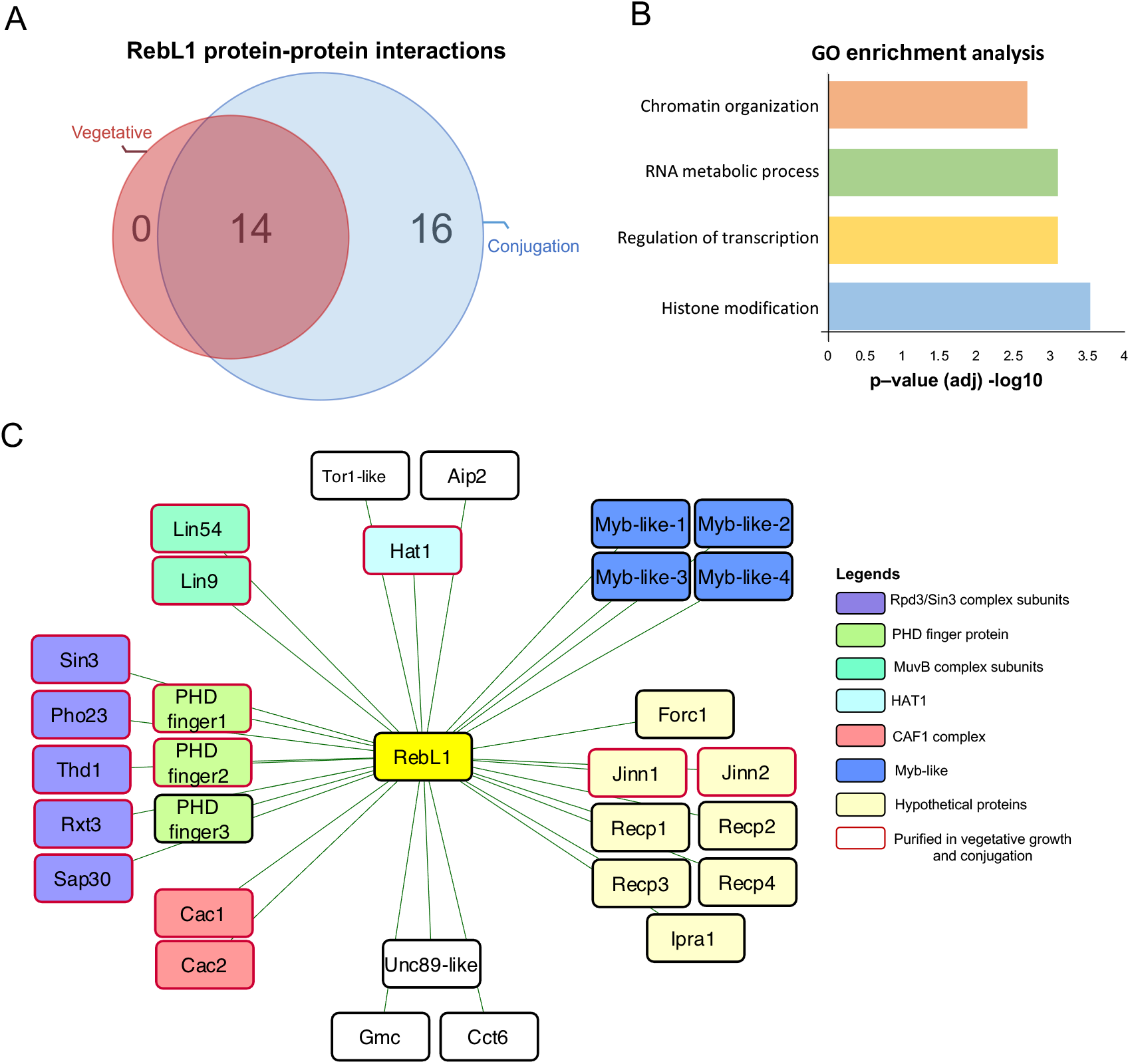
RebL1 has a conserved protein interaction network in *Tetrahymena*. **A:** Venn diagram illustrating the overlap of high confidence (FDR≤0.01) RebL1 co-purifying interaction partners during growth and conjugation (5h post-mixing the cells) in *Tetrahymena*. See Supplemental Tables S2-S3 for complete AP-MS results from vegetative and conjugating cells, respectively. **B:** GO enrichment analysis of RebL1-FZZ co-purifying proteins. **C:** Network view of RebL1-FZZ PPIs during conjugation (FDR≤0.01). Bait node is shown in yellow whereas the co-purifying partners are grouped according to their putative complexes.

Additionally, we identified TTHERM_00841280, which appears to be ciliate-specific and lacks sequence similarity to any known metazoan protein. The expression analysis of RebL1 co-purifying proteins using publicly available micro-array data revealed that TTHERM_00841280 is exclusively expressed during conjugation with a peak at early stages correlating with the onset of MIC meiosis (Supplemental figure S2) (77). Hence, we named TTHERM_00841280 as ‘***F***riend ***o***f ***R***ebL1 in ***c***onjugation’ (Forc1). Among the remaining conjugation-specific RebL1 co-purifying partners were a Tor1-like, Asf1-interacting protein 2 (Aip2) and several unannotated proteins, including Recp1-4 (***Re***bL1 ***c***o-purifying ***p***rotein 1-4) and Ipra1 (see below), without any recognizable domains (Figure 2C). We have previously shown that Aip1 and Aip2 are two highly similar ciliate-specific proteins that interact with Asf1 and likely function in chromatin assembly pathways in *Tetrahymena* (69). We also detected Aip1 as a conjugation-specific interaction partner of RebL1, albeit at a slightly relaxed threshold (FDR≤0.03) (Supplemental Table S3). The co-purification of Aip1/2 proteins that function with ImpB6 and Asf1 (69) reinforces the idea that RebL1 functions in multiple chromatin-related pathways in *Tetrahymena*, consistent with our GO enrichment analysis of the RebL1 interacting partners (Figure 2B).

### RebL1 protein interaction profile is analogous to human RBBP4 and RBBP7

To examine the functional conservation of distantly related orthologues, we directly compared the protein interaction profiles of RebL1 and human RBBP4 and RBBP7. We generated inducible HEK293 cell lines expressing EGFP-tagged variants of RBBP4 and RBBP7 and used them for anti-GFP AP-MS experiments. Using SAINTexpress, we found RBBP4 and RBBP7 to physically interact with multiple chromatin-related, as well as transcriptional regulatory, protein complexes (Supplemental Table S4,5), including the CAF1-, Hat1-, MuvB-, and Sin3A-complexes (FDR≤0.01; Figure 3A), as reported previously (78). However, 37.5% (18 proteins out of 48) of the high-confidence protein-protein interactions for RBBP4 and 29% (9 proteins out of 31) for RBBP7 had not been previously reported in the BioGRID database (79). While there are many shared interaction partners between the RBBP4 and RBBP7 homologues, non-overlapping distinct PPIs were also observed (Figure 3B). For example, consistent with previous studies (35), MuvB subunits co-purified with only RBBP4 and not with RBBP7 (Figure 3A). Furthermore, CAF1-complex subunits Chaf1A and Chaf1B co-purified with RBBP4 whereas Hat1 was identified exclusively in RBBP7 interaction partners. Importantly, we observed that many of the transcription-and chromatin-related complexes that we found for human RBBP4 and RBBP7 had their putative analogous complexes in *Tetrahymena*. Examples of these conserved interaction partners include the Hat1-, CAF1-, and MuvB (Figure 3C; Supplemental Table S3). Although, PRC2 subunits, including Suz12, Ezh2 and EED, were co-purified with RBBP4 and RBBP7 (Figure 3A), however, we did not identify any orthologues subunits in our RebL1 AP-MS data. Collectively, we conclude that *Tetrahymena* RebL1 physically interacts with multiple epigenetic regulatory complexes, and these functional links are conserved in humans.

**Figure 3:**
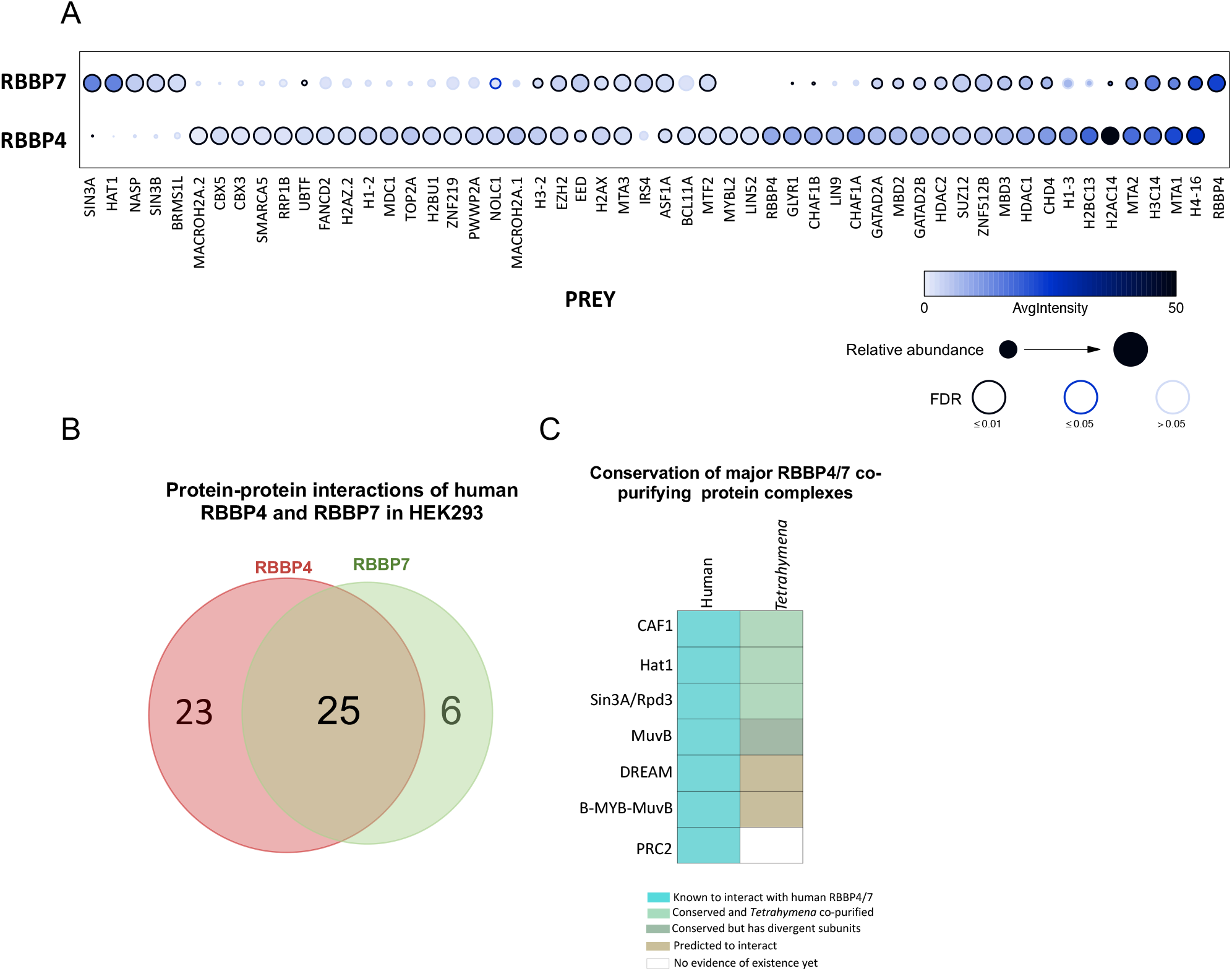
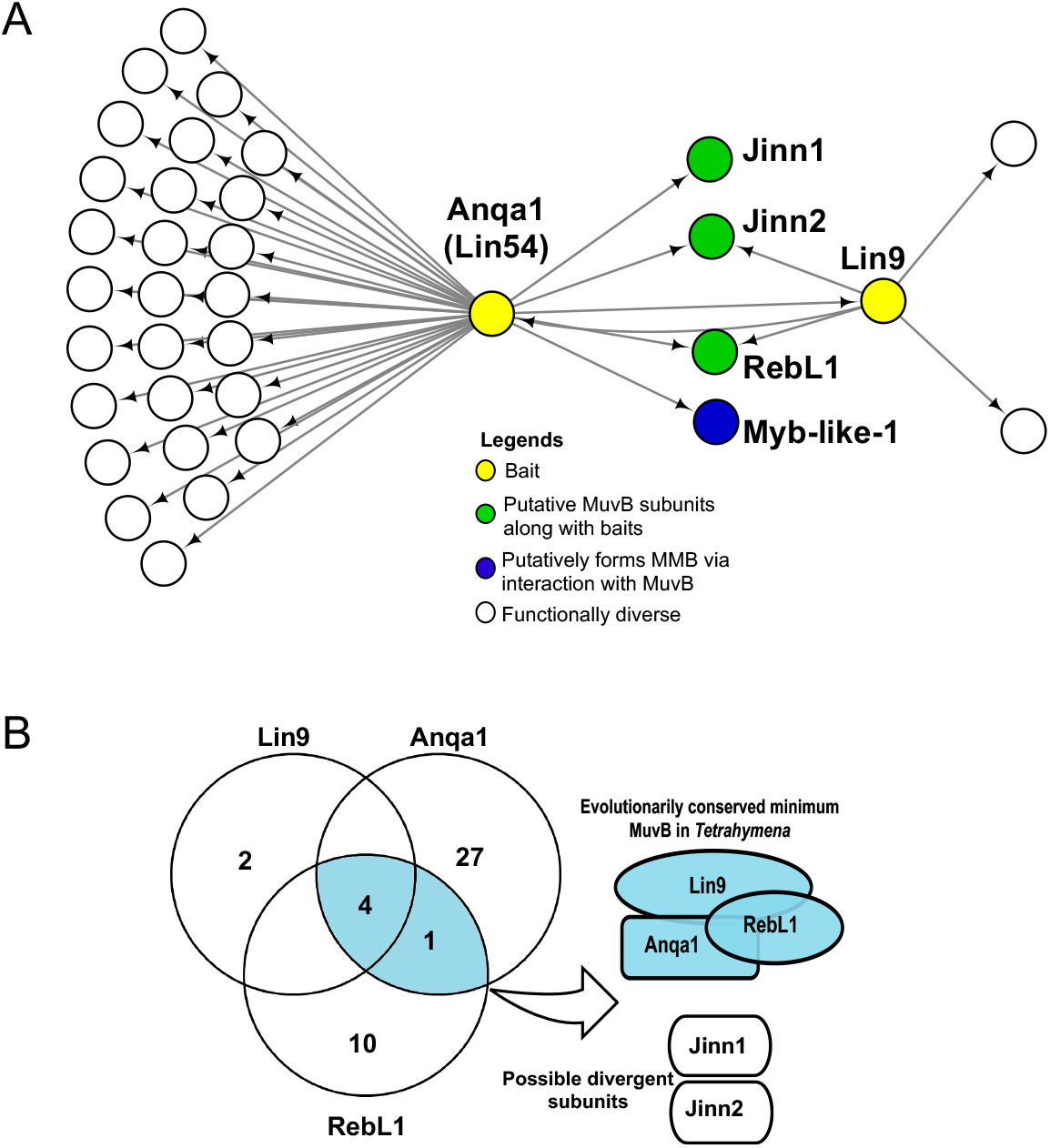
Proteomic analysis of human RBBP4 and RBBP7. **A:** Dot plot representation of the interaction partners identified with RBBP4 and RBBP7 in HEK293 cells. Inner circle color shows the average spectral count, the circle size indicates the relative prey abundance, and the circle outer edge is the SAINT FDR. **B:** Venn diagram of the overlap of PPIs for human RBBP4 and RBBP7 proteins (FDR≤0.01) from HEK293 cells. See Supplemental Tables S4-S5 for complete AP-MS results for RBBP4 and RBBP7, respectively. **C:** Schematic illustration of evolutionary conservation for the RebL1 co-purifying putative chromatin-related complexes that are also known to interact with human RBBP4 and RBBP7 proteins. Figure legend is provided. Figure 3: **Proteomic analysis of putative MuvB subunits in***Tetrahymena*. **A:** Network depiction of Lin9 and Anqa1 (Lin54) protein-protein interactions (FDR≤0.01). Bait nodes are shown in yellow. See Supplemental Tables S6-S7 for complete AP-MS results for Lin9 and Anqa1, respectively. **B:** Venn diagram indicating the overlapping PPIs of RebL1, Lin9 and Anqa1 during vegetative growth.

### RebL1, Lin9^Tt^ and Lin54^Tt^ are the conserved MuvB subunits

The identification of putative MuvB subunits in *Tetrahymena* piqued our interest due to the described role of mammalian MuvB as the master regulator of cell-cycle-dependent gene expression (33). To identify all the putative MuvB components, we initially used the human MuvB subunits as queries to search the *Tetrahymena* genome. Our analysis indicated that the *Tetrahymena* genome does not appear to encode any direct orthologues of Lin37 and Lin52. In addition, Lin9 appears to be present as a single gene copy in *Tetrahymena*, similar to humans. Remarkably, we identified at least 14 genes that appeared to encode the Lin54-like proteins in *Tetrahymena*. All the identified Lin54-like proteins contained two tandem TESMIN/TSO1-like CXC domains (Pfam: PF03638), consistent with the architecture of human Lin54 (39, 40)(Supplemental figure S3A). We next examined the expression profiles of the *Tetrahymena* Lin54-like genes during growth and conjugation using publicly available microarray data. Most of the Lin54-like genes were weakly expressed during the growth and late conjugation stages (6-hour and onwards after mixing the cells) (Supplemental figure S3B). At least 9 out of the 14 genes, including the RebL1 interaction partner TTHERM_00766460, exhibited an expression peak at early conjugation stages (2-6 hours after mixing the cells). For example, the expression of TTHERM_00120820, TTHERM_00577120 and TTHERM_00600840 peaked at 2 hours, whereas TTHERM_00766460 had its highest expression at 4 hours after mixing the cells (Supplemental figure S3B). These observations suggest that the expression of Lin54-like proteins is tightly regulated during early conjugation events, a period when the MIC becomes transcriptionally active and undergoes meiosis (Supplemental figure S1E). We named Lin54-like proteins as ‘Anqa-1 to 14’ (after the ‘ANQA’ mythological creature thought to appear once in ages), considering that several of them exhibited an expression peak only once at a distinct conjugation stage.

We next generated endogenously tagged Lin9^Tt^-FZZ and Anqa1-FZZ (TTHERM_00766460) cell lines and employed them for AP-MS (Supplemental figure S3C). These analyses showed reciprocal purification of RebL1 and Lin9^Tt^ when Anqa1-FZZ was employed as the bait protein (Figure 3A; Supplemental Tables S6,7). Similarly, Anqa1 and RebL1 were identified as Lin9^Tt^-FZZ high-confidence co-purifying partners. These results are consistent with the idea that RebL1, Anqa1 and Lin9^Tt^ represent the conserved subunits of a putative MuvB complex in *Tetrahymena*. Remarkably, two unannotated proteins, which we named as Jinn1 and 2 (see above), co-purified with both Lin9^Tt^-FZZ and Anqa1-FZZ as high-confidence hits (Figure 3A). Based on their expression profiles (Supplemental figure S2) and co-purification with all three conserved subunits (RebL1, Lin9^Tt^ and Anqa1) (Figure 3B), we suggest that Jinn1/2 represent *Tetrahymena*-specific putative MuvB components.

In addition to functionally diverse proteins such as kinases and helicases, TTHERM_00317260 and TTHERM_00842380 also co-purified with Anqa1-FZZ during vegetative growth (FDR≤0.01). TTHERM_00317260, which interacted with RebL1 during conjugation, is a *Tetrahymena*-specific protein and lacks any recognizable domains. We named it as Ipra1 (***I***nteraction ***p***artner of ***R***ebL1 and ***A***nqa1). TTHERM_00842380 is the putative MYB-like TF (MybL1) that was detected as a high-confidence RebL1 interaction partner during both growth and conjugation (FDR≤0.01) (Figure 3A). This observation is consistent with the reported interaction between mammalian MuvB and B-MYB TF to form an activator MMB complex (33). **Genome-wide analysis reveals distinct RebL1 and Anqa1 DNA-binding profiles**

Among the human MuvB subunits, RBBP4 and Lin54 are thought to bind DNA, with Lin54 exhibiting sequence specificity by recognising the CHR element (TTTGAA) (33). To investigate their genome-wide localization in *Tetrahymena*, we utilized our endogenously tagged RebL1 and Anqa1-FZZ cell lines and performed chromatin immunoprecipitation followed by high throughput sequencing (ChIP-seq) experiments. ChIP-seq analysis revealed that RebL1 peaks (5531 total peaks; q value < 0.05) predominantly localizes to genic regions (Figure 4A). The majority of the RebL1 peaks (64%) were found within exons, with only 2% being localized to intergenic regions (Figure 4A). Consistent with the role of RebL1 in multiple chromatin-related processes, metagene analysis indicated that RebL1 binds throughout the gene body of target genes without any specific sequence preferences (Figure 4B).

**Figure 4:**
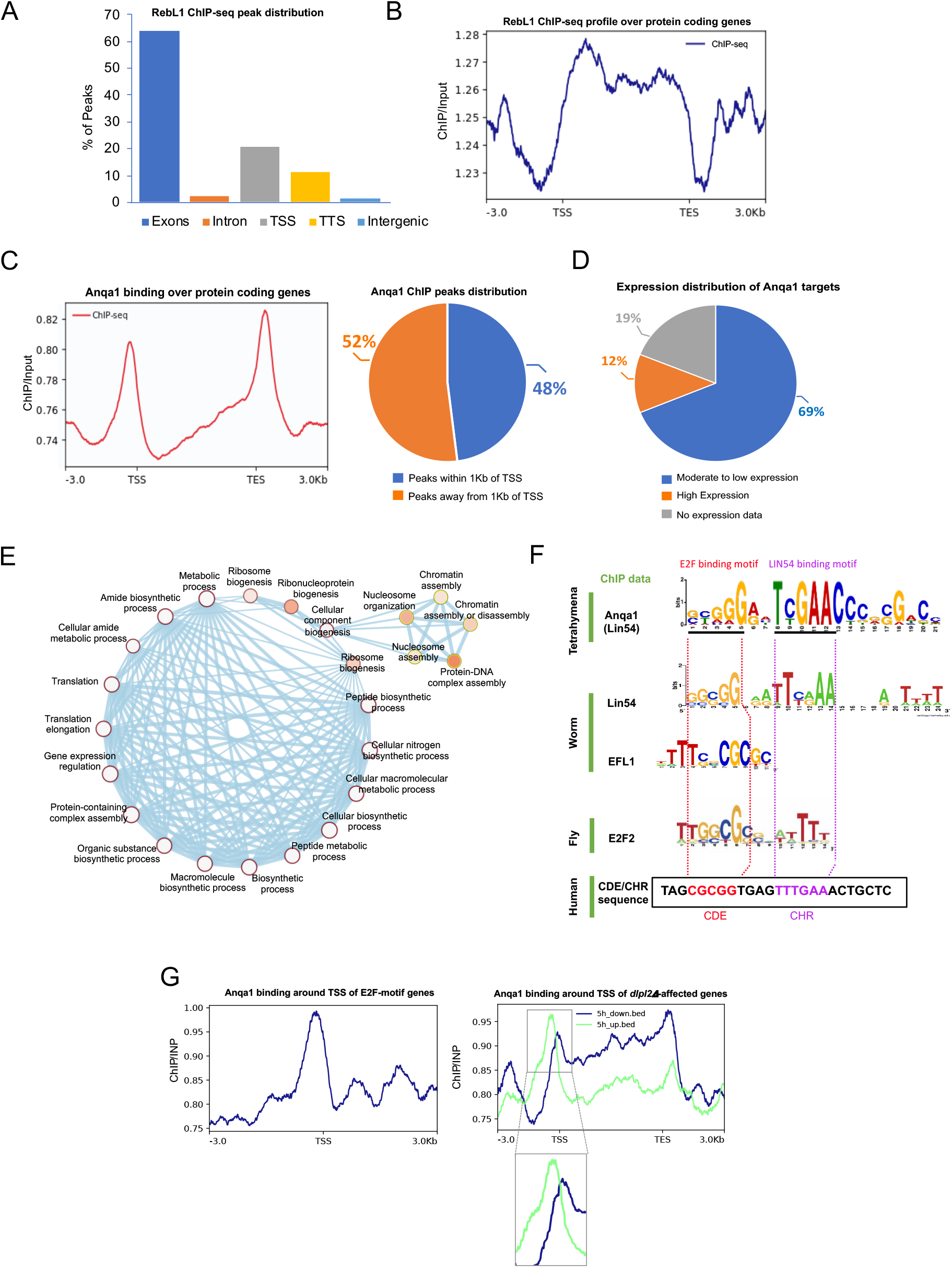
Genome-wide analysis of RebL1 and Anqa1 DNA-binding profiles in *Tetrahymena*. **A**: Bar plot of RebL1 ChIP peak distribution across various genomic features. **B:** Metagene analysis of RebL1 ChIP-seq data. **C:** Left: Metagene analysis of Anqa1 ChIP-seq data indicates enrichment near the TSS and TES of target genes. Right: Anqa1 ChIP peak distribution with respect to annotated TSS +1Kb. **D:** Pie chart showing the distribution of Anqa1 bound genes based on their expression levels. Anqa1 predominantly targets those genes that are moderately to very weakly expressed during vegetative growth. **E:** Network depiction of gene ontology (GO) enrichment analysis among Anqa1 bound targets. In this representation, each node depicts a GO term whereas each edge represents the degree of gene overlap that is found between two gene sets corresponding to the two GO terms. GO term significance 0.01 (white) to 0 (orange). **F:** An overrepresented motif in Anqa1 bound *Tetrahymena* genes identified by MEME (E value: 8.0e-223). Below are shown previously identified motifs for the *C. elegans* EFL-1, an extended motif enriched among E2F2, LIN-9 and LIN-54 co-regulated genes in *Drosophila* and the CDE/CHR motif from the human Cdc2 promoter. Regions bound by human E2F4 and LIN54 at the Cdc2 promoter as well as homologous motif sequences in other organisms are highlighted by dotted lines. **G:** Metagene analyses indicating the binding profile of Anqa1 around TSS of those genes that 1) contain the E2FL1 motif in their promoters (left), 2) those genes whose expression is affected during conjugation upon *DPL2* depletion (right).

Unlike RebL1, Anqa1 metagene analysis indicated a clear enrichment near the transcription start sites (TSS) and transcript end sites (TES) (Figure 4C). Consistently, roughly half of the Anqa1 ChIP peaks were identified within 1 kilobase of the annotated TSS (Figure 4C). As an enrichment near the promoter regions suggests a role in transcription regulation, we utilized publicly available RNA-seq data that has been used to rank *Tetrahymena* genes based on their expression patterns during vegetative growth (65) and correlated it with the Anqa1 ChIP data. We found that the majority of the Anqa1-FZZ bound genes (290 of 422 genes) are moderately to very weakly expressed, whereas only 12% of the targets are classified as highly expressed during *Tetrahymena* growth (Figure 4D). These results suggest that Anqa1 may function to regulate the expression of its target genes.

We next carried out Gene ontology (GO) enrichment analysis related to molecular functions and/or biological processes. While RebL1-binding genes were not enriched in any GO terms, in keeping with its role in multiple protein complexes, Anqa1 targets were however enriched for terms associated with gene expression regulation, chromatin assembly and organization, translation regulation, ribosome biogenesis and cellular metabolic processes, suggesting a specific function in regulation of cellular processes (Figure 4E).

### A hybrid E2F/DP and Anqa1 DNA-binding motif

The ChIP-seq analysis of Lin54 in metazoans has revealed the existence of a hybrid DREAM-recruiting DNA motif representing the E2F-binding sequence upstream of the CHR element (44– 46). To identify a putative DNA-binding motif, we performed a *de novo* DNA motif discovery analysis using the Anqa1 ChIP peaks. The identified Anqa1 putative motif appeared to share similarity with the characteristic DREAM-recruiting E2F/DP-CHR hybrid motif (Figure 4F). In *Tetrahymena*, at least 714 genes have been identified via *in silico* analysis to contain putative E2F DNA-binding motifs in their promoters (80). We found that Anqa1 exhibited enriched binding in the promoter regions of putative E2F motif containing genes (Figure 4G, left), consistent with the identification of an E2F/DP-CHR-like hybrid motif.

Although we did not detect any DREAM-specific proteins in our AP-MS (FDR≤0.01), the *Tetrahymena* genome does encode at least seven E2F family members (E2FL1-4 and DPL1-3) and two p107/p130 (Retinoblastoma-like1/2)-like proteins (81). In fact, an E2FL1 and Dpl2 heterodimer has been described to function in gene expression regulation during conjugation (81). To further explore the connection between Anqa1 and E2Fs, we utilized previously generated RNA-seq data in *dpl2Δ* cells during conjugation (81), and examined the Anqa1 binding profile around the TSS of affected genes. Our analyses indicated that Anqa1 binds relatively evenly across the gene bodies of both up- and down-regulated genes at 3,4,6 and 7h time points during conjugation (after mixing the *dpl2Δ* cells) (Supplemental figure S4). Remarkably, however, we observed a strong binding enrichment for Anqa1 specifically within the promoter regions of genes that are upregulated in *dpl2Δ* cells 5-hours post mixing cells (Figure 4G, right), a period in conjugation immediately prior to the observed meiotic defect in *dpl2Δ* cells (81). In contrast, the downregulated genes had wide-spread binding within their bodies without any enrichment near the TSS (Figure 4G, right). These *dpl2Δ*-dependent upregulated genes were enriched in pathways such as DNA replication and DNA repair (Supplemental figure S4). We conclude that the putative MuvB subunit Anqa1 binds in the promoter region, possibly in conjunction with DREAM-specific E2F/DP proteins, to regulate the expression of some of its target genes.

### RebL1 depletion results in growth defects

Previous studies have correlated the overexpression of RBBP4 and RBBP7 with tumor cell proliferation in certain cancer types (49, 54). To analyze the RBBP4 and RBBP7 expression levels and correlation with cancer prognosis, we utilized archived patient RNA-seq data from ‘The Cancer Genome Atlas’ (TCGA), focusing on multiple tumor types for which the adjacent normal tissue data were available. The expression levels of both RBBP4 and RBBP7 were significantly higher in several cancers, including stomach adenocarcinoma, breast cancer, glioblastoma multiforme (GBM), cholangio carcinoma and liver hepatocellular carcinoma (LIHC), compared with the normal tissues (p-value cut-off ≤0.01) (Supplemental figure S5A; Supplemental Tables S9 for TCGA abbreviations). Furthermore, these high expression levels of RBBP4 and RBBP7 significantly correlated with unfavorable clinical outcomes (P≤0.05) in multiple tumor types (Supplemental figure S5B). For example, Kaplan–Meier curves indicated that LIHC high expression groups for both RBBP4 and RBBP7 had significantly worse overall survival (P < 0.01) compared with low expression groups (Figure 5A). The observed overall poor survival of LIHC high-expression group was further enhanced when the RBBP4 and RBBP7 expression was combined in our analyses, suggesting an additive effect (Figure 5B).

**Figure 5:**
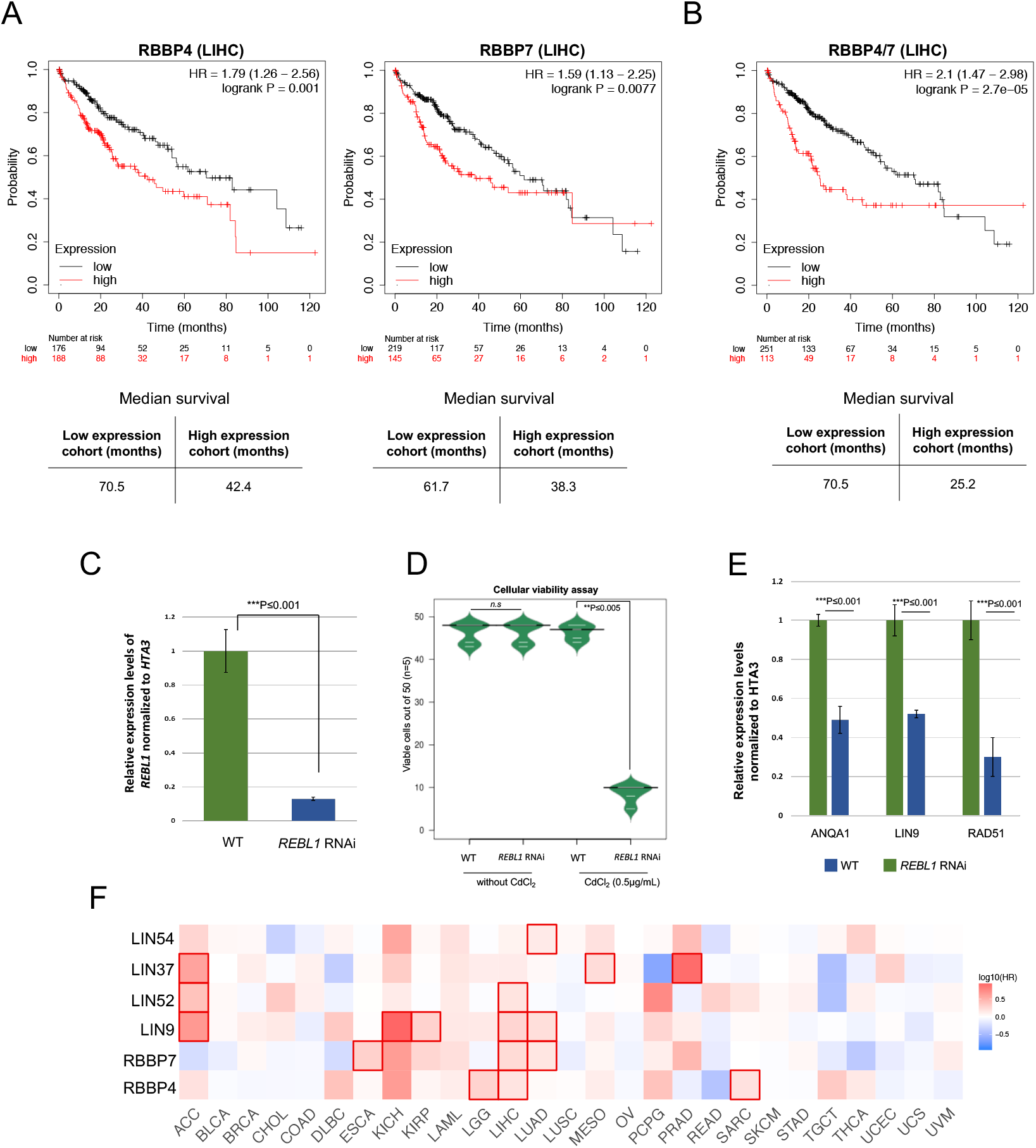
Conserved oncogenic properties of RBBP4/7. **A**: Kaplan–Meier curves of overall survival in patients with primary hepatocellular carcinoma using TCGA-LIHC. Patients were grouped according to median expression of either RBBP4 (left) or RBBP7 (right) using log-rank tests. Bottom panels show the median survival in months for high and low expression groups. **B:** Kaplan–Meier curves of overall survival in LIHC patients using combined expression of RBBP4 and RBBP7 to examine the survival rates in the TCGA-LIHC cohort. Log-rank test was used in this analysis and the bottom panel depicts the median survival in months for high and low expression groups. **C:** Expression levels of RebL1 in *RebL1* RNAi *Tetrahymena* cells treated with CdCl2 (0.5μg/mL) or not, as assessed by qRT-PCR. The qRT-PCR data were normalized to *HTA3* expression. ∗∗∗ indicates t-test p ≤ 0.001. The individual error bars indicate the standard deviation for each sample. **D:** Bean plot representation of cellular viability of single cells wild-type or *RebL1*-RNAi treated lines isolated in drops of media with and without CdCl2 (0.5μg/mL). Cellular growth was observed after at least 48 to a maximum of 72 hours. The number of viable drops were counted from each replicate (n=5; 50 cells in each replicate) and plotted. ∗∗ indicates student’s t-test p≤0.01. **E:** Analysis of expression levels of target genes in *RebL1*-RNAi cells treated with CdCl2 (0.075 μg/mL) or untreated. The qRT-PCR data were normalized to *HTA3* expression. ∗∗∗ indicates t test p ≤ 0.001. **F:** Heat map showing the hazard ratios in logarithmic scale (log10) for the indicated genes in multiple cancer types. The red blocks denote higher whereas blue blocks indicate lower clinical risk, with an increase in the gene expression. The blocks with darkened frames denote statistical significance in prognostic analyses (p≤0.05; Mantel–Cox test). The TCGA cancer abbreviations are provided in Supplemental Table S9.

We next examined the DepMap database, which encompasses data from genome-wide RNAi and CRISPR loss-of-function screens across hundreds of cancer cell lines, and observed that RBBP4 behaves as a common essential gene (Supplemental figure S5C) (82, 83). RBBP7, on the other hand, exhibited high selectivity in terms of cancer cell line dependency, reinforcing the concept of orthologue-specific functions for RBBP4 and RBBP7. Considering the sequence/structural similarity of RebL1 with its human orthologues and formation of similar functional links (Supplemental figure S5D), we explored the role of RBBP4 and 7 in cellular proliferation and/or gene expression using *Tetrahymena* as a model system. To this end, a cadmium inducible *RebL1* RNA interference (RNAi) construct was introduced into vegetative *Tetrahymena* cells. Upon cadmium chloride treatment, the RebL1 expression levels were significantly reduced in *RebL1*-RNAi cells in comparison with the WT and/or uninduced *Tetrahymena* (Figure 5C), as assessed by reverse-transcription quantitative real-time PCR (RT-qPCR) analysis. We used growth assays to examine the functional consequences of RebL1 depletion in growing *Tetrahymena* cells. We found that *RebL1*-RNAi cells gave significantly fewer (~19%) viable clones compared to wildtype *Tetrahymena* (90% viable cells; n=5) when treated with cadmium chloride (0.5μg/mL; see methods) (Figure 5D), consistent with the role of its human orthologues in cell proliferation.

### RebL1 depletion causes gene expression changes

The *RebL1*-RNAi dependent reduction in cellular viability could be related to indirect changes in expression levels of some functionally important genes. In human glioblastoma cells (GBMs), RBBP4 depletion results in sensitization of tumor cells to temozolomide (TMZ) in conjunction with the downregulation of DNA repair protein Rad51 (54). We examined Rad51^Tt^ expression levels in *REBL1*-RNAi cells. Rad51^Tt^ was significantly supressed in *Tetrahymena* depleted of RebL1 in comparison to the WT and/or uninduced cells (Figure 5E).

Previous studies have identified an auto-regulatory feedback loop for the DREAM complex (84). To test the possibility of an auto-regulatory feedback loop for MuvB, we examined the expression levels of Anqa1 and Lin9^Tt^ using the RNAs derived from either the *REBL1-*RNAi or WT cells. We found that *REBL1* knockdown suppressed the expression of both Anqa1 and Lin9^Tt^ (Figure 5E), consistent with the disruption of a putative *Tetrahymena* MuvB. Consistent with these data, we observed that expression levels of Lin54 and Lin9 positively correlate with RBBP4 in multiple tumor types (Supplemental figure S6). Furthermore, the high expression of MuvB subunits correlated with significantly worse overall survival in many different cancer types (P≤0.05), particularly in LIHC patients where increased levels of 3 out of the 5 MuvB subunits were associated with unfavourable outcomes (Figure 6C). These results are consistent with the highly conserved physical interactions of MuvB subunits with each other.

## Discussion

Effective transcriptional regulation requires the ordered assembly of multiple protein complexes on chromatin in response to specific cellular cues. In this study, we report that *Tetrahymena* contains at least three WD40 repeat family proteins with putative scaffolding roles. Specifically, we determined TTHERM_00688660 (RebL1) to be the major structurally and functionally conserved orthologue of human RBBP4 and 7 in *Tetrahymena*. We demonstrated that RebL1 is a component of multiple chromatin-related complexes and is important for gene expression regulation and cellular growth.

Previous studies have indicated that the structure/function of various MuvB-derived complexes can differ across different species (33). For example, structurally and functionally distinct MuvB complexes in *Drosophila* salivary glands and testis have been identified (85–87). It has remained an open question, however, whether tissue-specific or development-specific MuvB complexes also exist in other species. In *Tetrahymena*, the presence of multiple Lin54 paralogues and their distinct expression pattern suggest the existence of multiple MuvB-like complexes. Although further studies will explore this interesting possibility, a previous report has shown the interaction between E2FL1/Dpl2 and Anqa14 (81), supporting the existence of multiple structurally/functionally distinct MuvB-containing DREAM-like complexes in *Tetrahymena*. Several observations reported in our study are indirectly support the formation of a DREAM-like complex in *Tetrahymena*: 1) the identification of a putative E2F/DP-CHR-like DNA motif; 2) the significant enrichment of Anqa1 ChIP peaks in the promoter regions of putative E2F target genes in *Tetrahymena*; and 3) the targeting of *dpl2Δ-*dependent upregulated genes by Anqa1. We propose that *Tetrahymena* MuvB functions in collaboration with E2F/DPs to form a DREAM-like complex that negatively regulates the expression of target genes. Although E2F DNA-binding motifs (consensus TTTSSCGC, where S is either a G or a C) are largely conserved during evolution (88), CHR elements have not yet been studied in *Tetrahymena*. Further studies are warranted to thoroughly examine the role of CHR and other *cis* elements in *Tetrahymena* cell cycle regulation.

The role of RBBP4 and 7 in cellular proliferation has been widely reported, although the mechanistic details have remained unclear. Our results suggest that one of the mechanisms through which RBBP4 and 7 might function in cellular growth is via regulating the cell-cycle regulatory complex MuvB. In *Tetrahymena* MuvB subunits appear to regulate their own expression levels, suggesting an autoregulatory feedback mechanism. Consistent with this, we found that the expression profiles of MuvB subunits are also interconnected in tumor cells. In previous reports, positive autoregulatory feedback loops have been reported involving B-Myb and the DREAM complex (84). It will be interesting to examine whether B-Myb also functions in the MuvB auto-regulation pathway. Considering that MuvB subunits were identified as biomarkers of poor prognosis for multiple cancers, our mechanistic findings have direct implications for better understanding tumor cell proliferation.

RBBP4 and 7 function in multiple epigenetic complexes which might have a role in cellular proliferation independent of MuvB. For example, knockdown of RBBP4 in GBM tumor cells results in suppression of the DNA-repair protein Rad51 due to disruption of the RBBP4/p300/CBP chromatin-modifying complex (54). Loss of RBBP4 in GBMs was found to be associated with suppression of tumor growth (54). Consistently, we observed that RebL1 depletion resulted in suppression of the Rad51^Tt^ orthologue in *Tetrahymena*. Disruption of *Tetrahymena RAD51^Tt^* has been reported to lead to severe abnormalities during vegetative growth, as well as in conjugation (89, 90). It remains to be seen whether Rad51^Tt^ also has a role in the reduction of cellular viability observed in *REBL1*-RNAi treated *Tetrahymena* cells. In summary, our research has extended the current understanding of transcription regulation in ciliates and, more broadly, on the multifaceted role (s) of RBBP4 and RBBP7 proteins in eukaryotes.

## Material and Methods

### Cell strains

*Tetrahymena* strains CU428 [Mpr/Mpr (VII, mp-s)] and B2086 [Mpr+/Mpr+ (II, mp-s)] of inbreeding line B were acquired from the *Tetrahymena* Stock Center, Cornell University, Ithaca N.Y. (http://tetrahymena.vet.cornell.edu/). Cells cultured in 1×SPP were maintained axenically at 30 °C as previously described (69). HEK293 cells (Flp-In 293 T-REx cell lines) were obtained from Life Technologies (R780-07) (Invitrogen) and cells were cultured in Dulbecco’s modified Eagle’s medium with 10% FBS and antibiotics as described (91).

### Macronuclear gene replacement

Epitope tagging vectors for histone HHF1, HHF2, RebL1, Anqa1 and Lin9^Tt^ were constructed as previously described (69). Two separate ~ 1 kb fragments up and downstream of the predicted stop codons were amplified using wild-type *T. thermophila* genomic DNA as template and primers as indicated in Supplemental Table S8. The resulting PCR products were digested with KpnI and XhoI (upstream product) or NotI and SacI (downstream product). The digested products were cloned into the tagging vector (pBKS-FZZ) kindly provided by Dr. Kathleen Collins (University of California, Berkeley, CA). The final plasmid was digested with KpnI and SacI prior to transformation. One micrometer gold particles (60 mg/ml; Bio-Rad) were coated with at least 5μg of the digested plasmid DNA. The DNA coated gold particles were introduced into the *T. thermophila* MAC using biolistic transformation with a PDS-1000/He Biolistic particle delivery system (Bio-Rad). The transformants were selected using paromomycin (60μg/ml). MAC homozygosity was achieved by growing the cells in increasing concentrations of paromomycin to a final concentration of 1 mg/ml.

### Epitope tagging in HEK293 cells

We cloned gateway-compatible entry clones for RBBP4 and RBBP7 into the pDEST pcDNA5/FRT/TO-eGFP vector according to the manufacturer’s instructions. The cloned vectors were co-transfected into Flp-In T-REx 293 cells together with the pOG44 Flp recombinase expression plasmid. Cells were selected for FRT site-specific recombination into the genome and stable Flp-In 293 T-REx cell lines were maintained with hygromycin (Life Technologies, 10687010) at 2ug/ml. Expression of the gene of interest was induced by addition of doxycycline to the culture medium 24 h before harvesting.

### Experimental design for mass spectrometry

We processed independently at least two biological replicates of each bait along with negative controls in each batch of sample. *Tetrahymena* cells without tagged bait (i.e. empty cells) and HEK293 cells expressing GFP alone were used as controls in our analysis. We performed extensive washes between each sample (see details for each instrumentation type) to minimize carry-over. Furthermore, the order of sample acquisition on the mass spectrometer was reversed for the second replicate to avoid systematic bias. On the LTQ mass spectrometer, a freshly made column was used for each sample as previously described (69).

### Affinity purification and Mass Spectrometry sample preparation

For *Tetrahymena* samples, affinity purification was carried out essentially as described (69). Briefly, *T. thermophila* was grown in ~500ml of 1×SPP to a final concentration of 3×10^5^ cells/ml. The cells were grown to mid-log phase for vegetative samples. For conjugation, cells were pelleted at 5h post mixing the different mating types of *Tetrahymena*. The cells were pelleted and frozen at −80C. The frozen pellets were thawed on ice and re-suspended in lysis buffer (10 mM Tris–HCl (pH 7.5), 1 mM MgCl_2_, 300 mM NaCl and 0.2% NP40 plus yeast protease inhibitors (Sigma)). For nuclease treatment, 500 units of Benzonase (Sigma E8263) was added and extracts were rotated on a Nutator for 30 min at 4°C. WCEs were clarified by centrifugation at 16,000×g for 30 minutes. The resulting soluble material was incubated with 50μL of packed M2-agarose (Sigma) at 4°C for at least 2 hours. The M2-agarose was washed once with 10 mL IPP300 (10 mM Tris– HCl pH 8.0, 300 mM NaCl, 0.1% NP40), twice with 5mL of IP100 buffer (10 mM Tris–HCl pH 8.0, 100 mM NaCl, 0.1% NP40), and twice with 5mL of IP100 buffer without detergent (10 mM Tris–HCl pH 8.0, 100 mM NaCl). 500uL of 0.5M NH_4_OH was added to elute the proteins by rotating for 20 minutes at room temperature. Preparation of protein eluates for mass spectrometry acquisition is detailed in supplementary methods.

### AP-MS procedure in HEK293 cells

The AP-MS procedure for HEK293 cells was performed essentially as previously described (91– 93). In brief, ∼20×10^6^ cells were grown independently in two batches representing two biological replicates. Cells were harvested after 24 h induction of protein expression using doxycycline. HEK293 cell pellets were lysed in high-salt NP-40 lysis buffer (10 mM Tris-HCl pH 8.0, 420 mM NaCl, 0.1% NP-40, plus protease/phosphatase inhibitors) with three freeze-thaw cycles. The lysate was sonicated as described (91) and treated with Benzonase for 30 min at 4⁰C with end-to-end rotation. The WCE was centrifuged to pellet any cellular debris. We immunoprecipitated GFP-tagged RBBP4 and RBBP7 proteins with anti-GFP antibody (G10362, Life Technologies) overnight followed by a 2-hour incubation with Protein G Dynabeads (Invitrogen). The beads were washed 3 times with buffer (10mM TRIS-HCl, pH7.9, 420mM NaCl, 0.1% NP-40) and twice with buffer without detergent (10mM TRIS-HCl, pH7.9, 420mM NaCl). The immunoprecipitated proteins were eluted with NH4OH and lyophilized. Proteins for MS analysis were prepared by in-solution trypsin digestion. Briefly, we resuspended the protein pellet in 44uL of 50mM NH_4_HCO_3_, reduced the sample with 100mM TCEP-HCL, alkylated with 500mM iodoacetamide, and digested it with 1ug of trypsin overnight at 37°C. Samples were desalted using ZipTip Pipette tips (EMD Millipore) via standard procedures. The desalted samples were analyzed with an LTQ-Orbitrap Velos mass spectrometer (ThermoFisher Scientific). Raw MS data were searched with Maxquant (v.1.6.6.0)(94) as described previously (93), yielding spectral counts and MS intensities for each identified protein for individual human proteins for each experiment. The resulting data were filtered using SAINTexpress to obtain confidence values utilizing our two biological replicates. The AP-MS data generated from HEK293 cells expressing GFP alone were used as negative control for SAINT analysis.

### MS data visualization and archiving

We used Cytoscape (V3.4.0; (95)) to generate protein-protein interaction networks. For better illustration, individual nodes were manually arranged in physical complexes. Dot plots and heatmaps were generated using ProHits-viz (96). The annotation of the co-purifying partners was carried out using BLAST (https://blast.ncbi.nlm.nih.gov/Blast.cgi). SMART domain analysis (http://smart.embl-heidelberg.de/) of the predicted proteins was carried out. All MS files used in this study were deposited at MassIVE (http://massive.ucsd.edu). Additional details (including MassIVE accession numbers and FTP download links) can be found in Supplemental Table S10. The password to access the MS files prior to publication is……

### ChIP-Seq and related analyses

The ChIP experiments were performed as described previously and see supplemental methods for details (70, 97). The ChIP-seq reads were mapped to *T. thermophila* genome (2014 annotation) using RACS pipeline essentially as reported (98). ChIP peaks were called using MACS2 software where the peak’s q-value must be lower than 0.05 (99). The ChIP inputs were used as the control. Subsequently, the identified summits (apex of the peak) with 50 bp extension in both directions were analyzed using MEME-ChIP software for *de novo* motif discovery (http://meme-suite.org/) (100). The metagene analysis was performed using ChIP-Seq reads normalized over the inputs and by ‘Reads Per Kilobase of transcript per Million mapped reads (RPKM)’ values. The plots were generated using deeptools (101).

### Gene expression data

For gene expression analysis, we used microarray data (accession number GSE11300) (http://tfgd.ihb.ac.cn/) (77) and the expression values were represented in the heatmap format. To assess the similarities in gene expression profiles hierarchical clustering was performed. RNA-seq data corresponding to *DPL2* knockout was acquired from GSE104524 and analyzed for differential expression as described previously (81).

### RNAi vector construction and RT-qPCR experiments

RNAi vectors were constructed essentially as described (102). Briefly, RebL1 target sequence used in hairpin RNA constructs was amplified from CU428 genomic DNA using PrimeSTAR Max DNA polymerase with primers as listed in Supplemental Table S8. To create the hairpin cassette, amplified target fragments were cloned into the BamHI–BamHI and PstI–PstI sites, respectively, of pAkRNAi-NEO5, which was kindly provided by Dr. Takahiko Akematsu, with the NEBuilder HF DNA Assembly kit. In this system, the NEO5 cassette of the backbone plasmid, which confers paromomycin resistance, has been replaced by a puromycin resistance marker (PAC) under the MTT2 copper-inducible promoter (102). The resulting plasmid was linearized with SacI and KpnI before biolistic transformation. The PAC cassette was activated by adding 630 μM CuSO4 to the cells, with the addition of 200 μg/mL puromycin dihydrochloride (Cayman Chemical, Ann Arbor, MI, USA, Cat. CAYM13884-500) to select transformants. Finally, we induced RNAi in cells carrying the hairpin construct by the addition of 0.5μg/mL CdCl_2_ during vegetative growth (1μg/mL CdCl_2_ was slightly toxic in WT cells). The expression of the target gene was examined using Rt-qPCR analysis. To this end, we isolated total RNA from the RNAi-treated or WT *Tetrahymena* cells using TRIzol (Life Technologies) as per the supplier’s instructions. The isolated RNA was treated with Deoxyribonuclease I (RNase-free, Thermo). cDNA was prepared using iScript™ Reverse Transcription Supermix for RT-qPCR. qPCR was performed in technical triplicates using the cDNA prepared from three individual KD cell lines. The data were normalized to the expression levels of *HTA3*. Primers are indicated in Supplemental table S8.

### Viability assays

Viability assays were carried out essentially as described by (103). We initiated our analysis by isolating single cells of wild-type or *RebL1*-RNAi treated lines in drops of media with and without CdCl2 (0.5μg/mL; note: 1μg/mL CdCl2 was slightly toxic in WT cells). After 2 days of growth, the approximate number of cells in each drop was determined and drops containing >1000 cells were considered as viable. The experiment was conducted in five independent biological replicates with 50 cells isolated in each replicate. Finally, the number of viable drops from each replicate for each condition was represented as a bean plot. We used student’s t-test to assess any statistically significant difference in the averages of viable drops of examined conditions.

### Data Deposition

The mass spectrometry data reported in this paper are at (https://massive.ucsd.edu/). Additional files include the complete SAINTexpress outputs for each dataset as well as a ‘‘README’’ file that describes the details of mass spectrometry files deposition to the MassIVE repository. ChIP-seq data generated can be found online at Gene Expression Omnibus (GEO, https://www.ncbi.nlm.nih.gov/geo/). NGS and reads files produced in this study were deposited at GEO database with unique identifier----.

## Acknowledgements

We thank Anita Samardzic for her technical assistance with *Tetrahymena* growth media preparations. We thank Dr. Ernest Radovani, Guoqing Zhong and Hongbo Guo for their technical help with HEK293 MS experiments. Shuye Pu is acknowledged for advice on ChIP analysis. The authors also wish to thank Dr. Anne-Claude Gingras at the Network Biology Collaborative Centre, Lunenfeld-Tannenbaum Research Institute, Mt. Sinai Hospital, Toronto, ON, Canada for access to mass spectrometers regarding *Tetrahymena* work. Tanja Durbic and team members at the Donnelly Sequencing Centre are gratefully acknowledged for their assistance with next-generation sequencing.

## Authors’ contribution

S.N.-S. and J.F jointly conceived the project. S.N.-S. generated cell lines, performed Western blots and affinity purifications, conducted TCGA analysis, ChIP data analysis and evolutionary studies, and wrote the manuscript. J.Garg generated cell lines, participated in affinity purifications, performed IF microscopy and RNAi experiments with contribution from J.Fine. AS performed ChIP-seq and participated in affinity purifications with contribution from SW and JD. KA generated cell lines, performed affinity purifications, and participated in TCGA analysis. HL and MP carried out ChIP data analysis. NA performed qPCR experiments and participated in figure preparation. SP and EM contributed to generation of HEK293 cell lines and related MS analysis. ZZ supervised HL. JG edited the manuscript, participated in analyses, and provided reagents. RP edited the manuscript, supervised trainees, and provided reagents. J.-P.L. performed all *Tetrahymena* MS analyses, contributed to data analysis and experimental design, partly supervised the project, and edited the manuscript. J.F. was responsible for study design, manuscript editing, data analyses, and project supervision.

All authors have approved the final manuscript.

