## Supplemental Figures 1-6 for "Functional characterization of RebL1 highlights the evolutionary conservation of oncogenic activities of the RBBP4/7 orthologue in *Tetrahymena thermophila*"

Nabeel-Shah_Figure S1

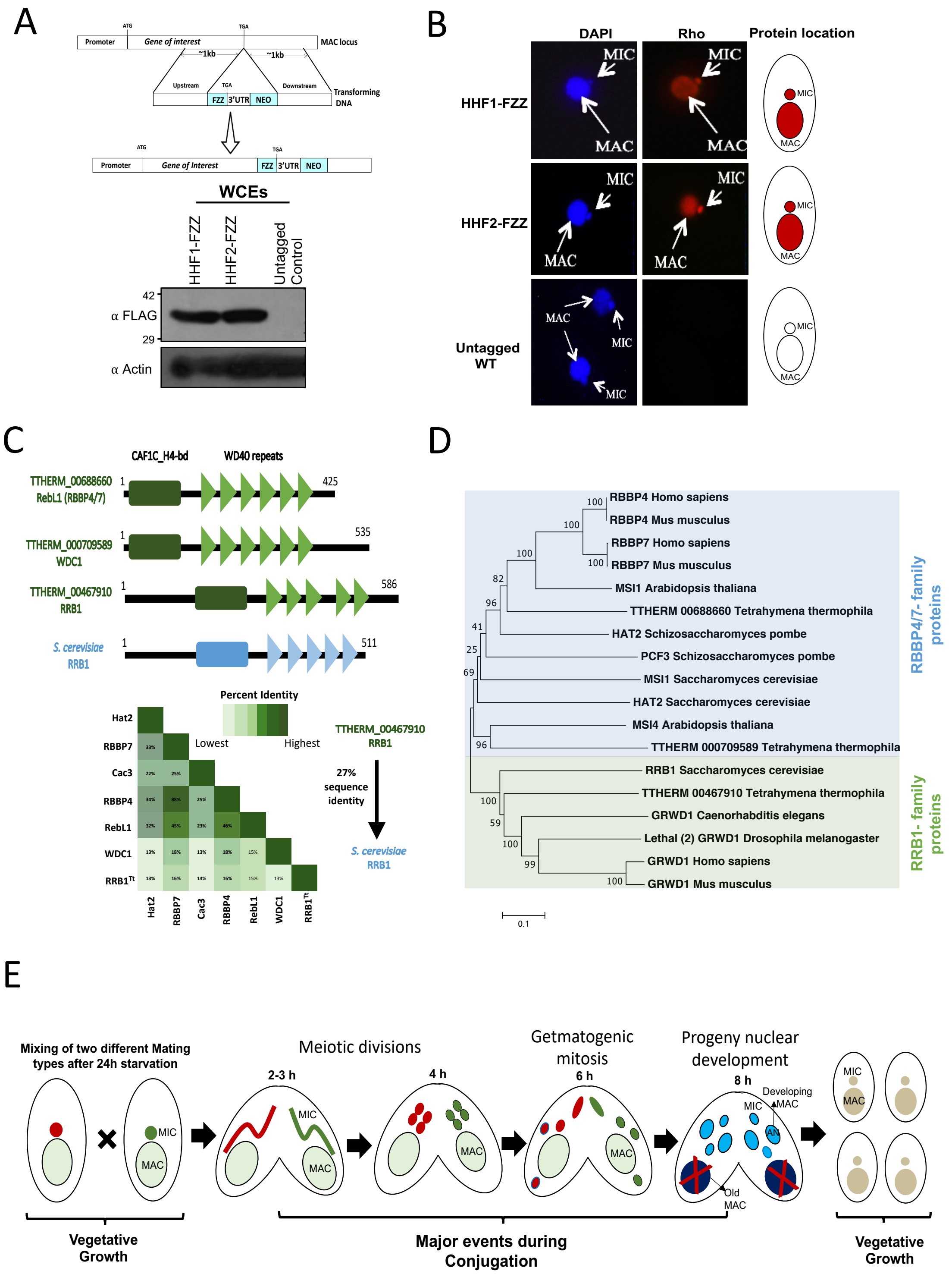

**Figure S1:** Characterization of RBBP4/7$^{Tt}$ in *Tetrahymena*. **A A:** Top: Schematic illustration of epitope tagging strategy for the MAC locus. Bottom: Western blotting analysis using whole cell lysates prepared from HHF1-FZZ and HH2-FZZ expressing *Tetrahymena* cells. The blot was probed with the indicated antibodies. **B:** Indirect Immunofluorescence (IF) analysis of H4-FZZ during vegetative growth in *Tetrahymena*. For nuclear counterstaining DAPI was used. MAC and MIC position are indicated. Cartoon diagram of the localization pattern is shown on the right. **C:** Top: Comparative domain analysis of RBBP4/7$^{Tt}$ (RebL1) protein against two other WD-40 repeat proteins in *Tetrahymena* and budding yeast RRB1. Bottom: Percent sequence identity shown as half-square rectangle. **D:** Neighbour-joining phylogenetic analysis of WD40 proteins. Different subfamilies are highlighted in different colors. The numbers on the branches represent confidence values based on 1000 bootstrap replicates. **E:** Life cycle of *Tetrahymena*. Parental macronuclei are shown in light green. After pairing of two cells of different mating types, the respective MICs (shown in red and dark green) undergo meiosis and adopt a crescent shape. One of the four meiotic products is selected in each cell and the other three are degraded. The selected meiotic product nuclei in each cell undergo mitosis and identical pro-nuclei are produced. One of two pro-nuclei in each cell is transferred to the mating partner. The exchanged nuclei fuse with the other resident pronucleus giving rise to a zygotic diploid nucleus that divides mitotically twice to generate four nuclei. Two of these resultant nuclei will develop as new MACs (anlagen), and two will become MICs. The parental MAC degenerates. Time periods shown correspond to hours post mixing of the two cells. The AP-MS for RebL1-FZZ was performed at 5h post mixing.

Nabeel-Shah_Figure S2

A

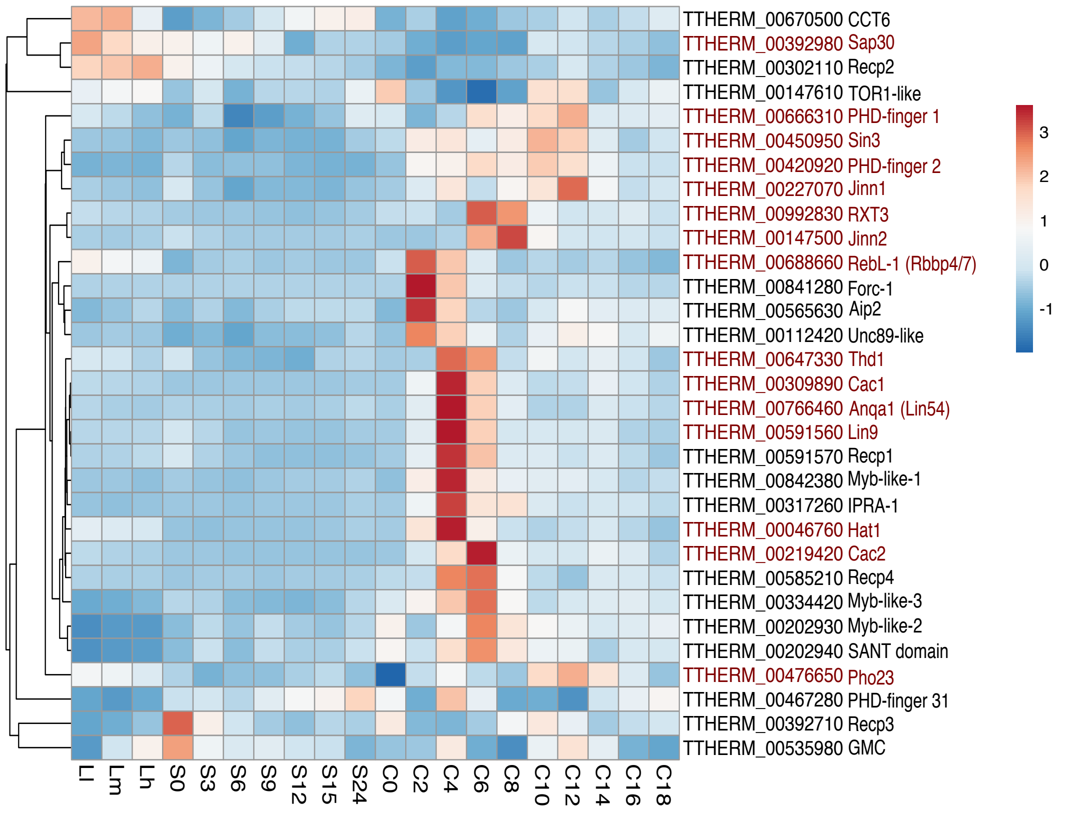


B

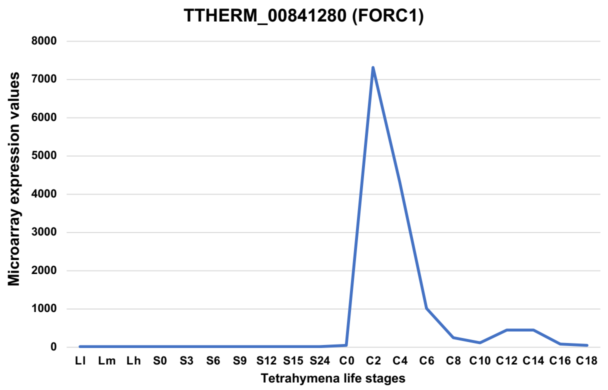

**Figure S2:** Expression analysis of RebL1-FZZ co-purifying proteins in *Tetrahymena*. **A:** Heat map display of microarray expression values for RebL1-FZZ co-purifying proteins. Z-scores were calculated across the rows for each gene to examine its differential expression across growth, starvation, and developmental stages. L1–LH display logarithmic growth phase, S0–24 indicates starvation for 24 h, and C is for conjugation where 0–18 are hours post mixing the different mating types. Hierarchical clustering was performed to assess the similarities in expression profiles. The labels in red are those proteins that were detected in both conjugation and vegetative growth of *Tetrahymena*. **B:** Line graph depiction of FORC1 expression which was co-purified with RebL1-FZZ during conjugation and is exclusively expressed during conjugation/development.

Nabeel-Shah_Figure S3

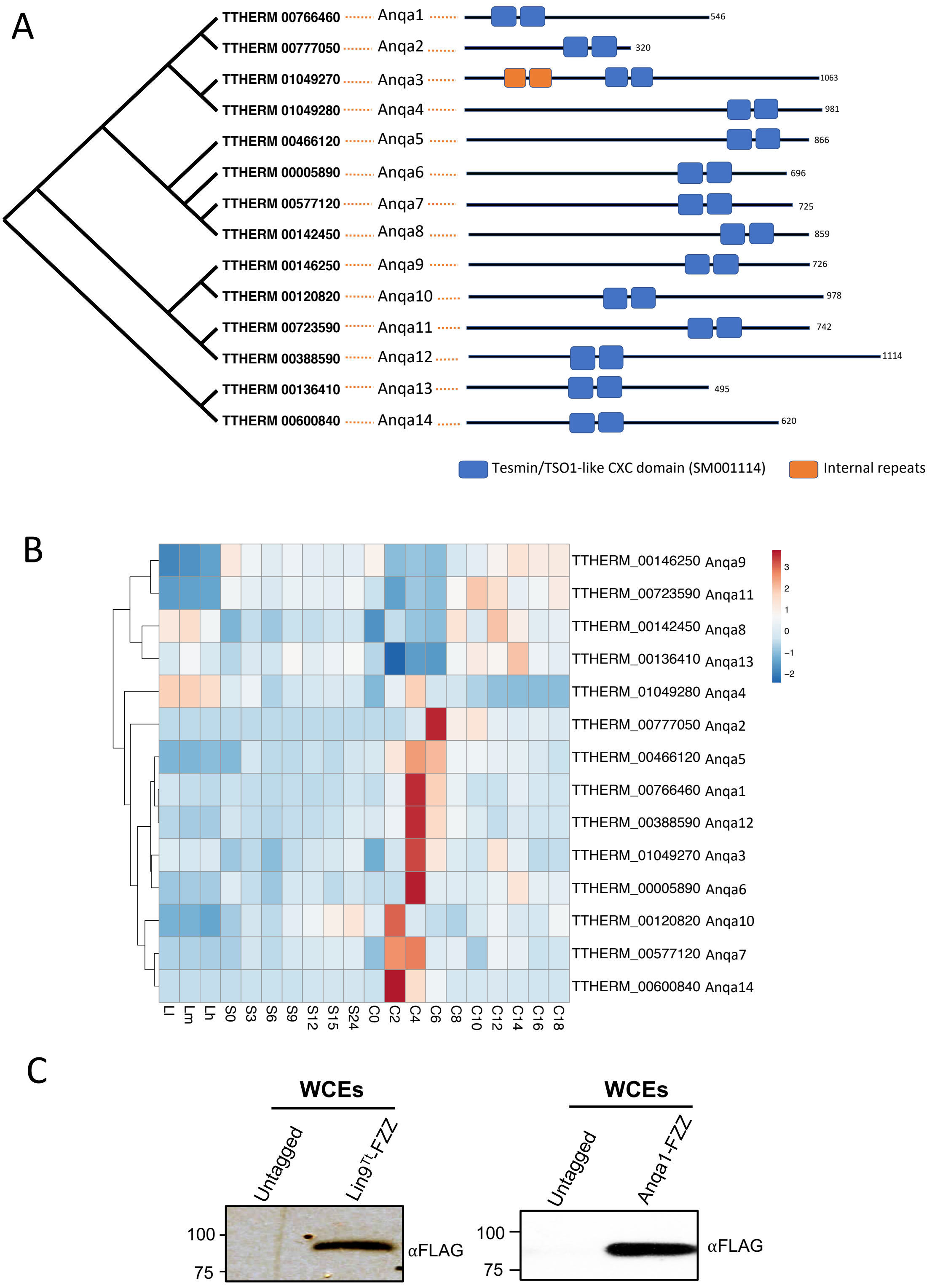

**Figure S3: A:** Neighbor-joining phylogenetic tree of Lin54-like proteins in *Tetrahymena*. The tree topology represents the bootstrap consensus tree (1000 bootstrap replicates) and branch lengths do not indicate the genetic distances. On the right side of the phylogenetic tree is the predicted domain architecture of each protein as well as their systematic nomenclature. Color code indicated. **B:** Heat map display of microarray expression values for Lin54-like genes in *Tetrahymena*. Z-scores were calculated across the rows for each gene to examine its differential expression across growth, starvation, and developmental stages. L1–LH: growth phase, S0–24: starvation for 24 h, and C conjugation where 0–18 are hours post mixing the different mating types. Hierarchical clustering was performed to assess the similarities in expression profiles. **C:** Western blotting analysis using whole cell lysates prepared from Lin9-FZZ (left) and Anqa1-FZZ (right) expressing *Tetrahymena* cells. The blot was probed with the indicated antibodies.

Nabeel-Shah_Figure S4

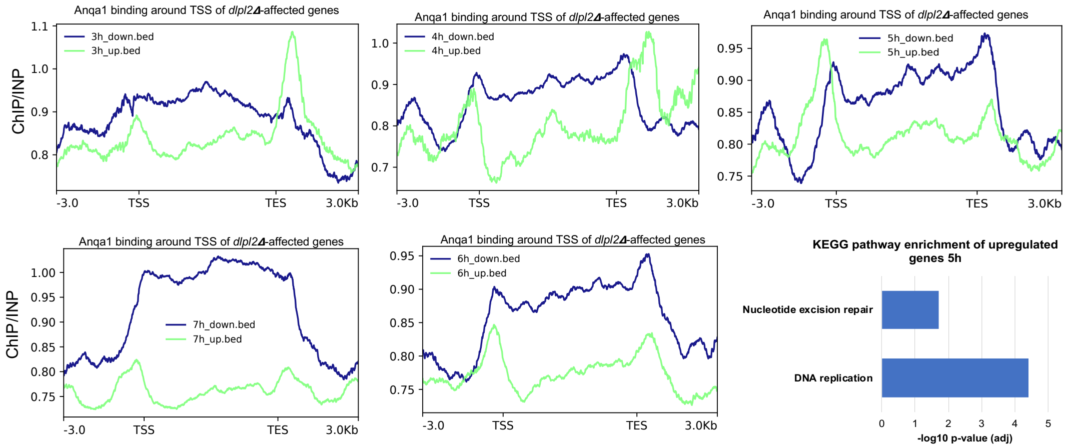

**Figure S4:** Metagene analyses indicating the binding profile of Anqa1 around TSS of those genes whose expression is affected during conjugation (conjugation time points 3h,4h,5h,6h, and 7h post mixing the cells) upon *DPL2* depletion (acquired from public database GSE104524). Blue and Green show the Anqa1 binding for downregulated and upregulated genes, respectively. Below the 5h plot is shown the KEGG pathway enrichment analysis for the upregulated genes which were preferentially bound by Anqa1 in the promoter region.

A

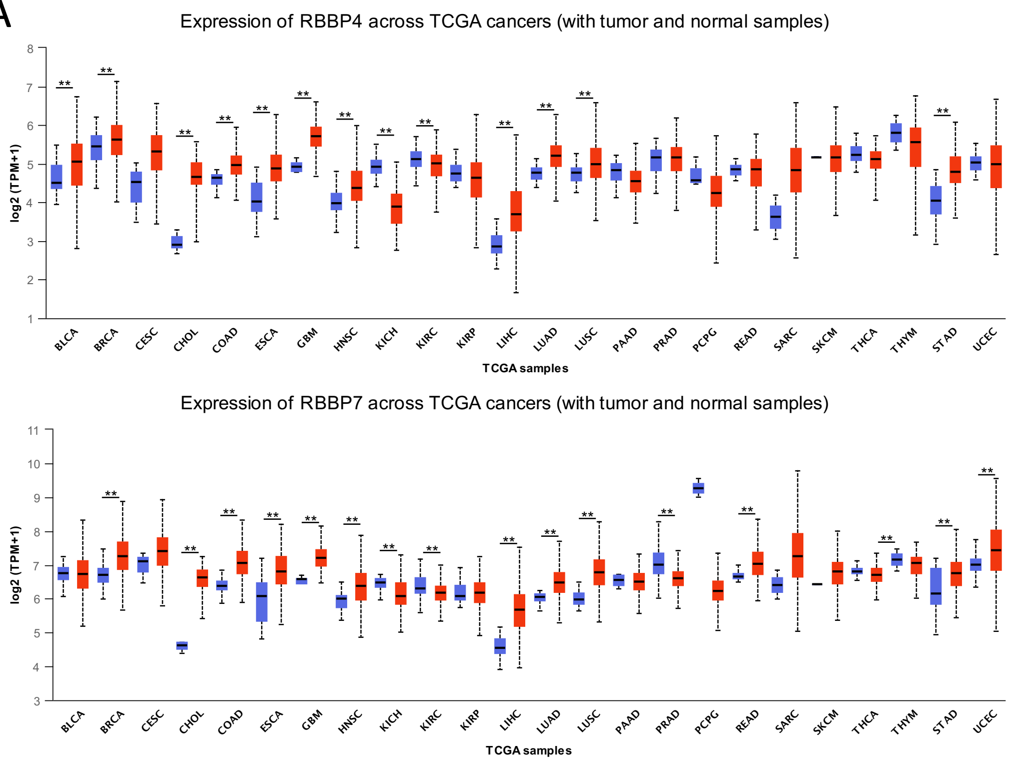


B

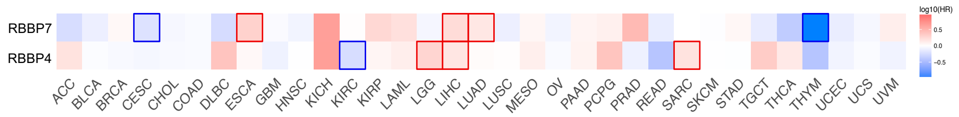


C

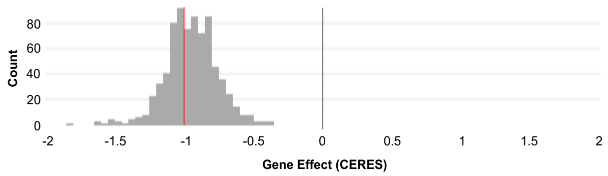


D

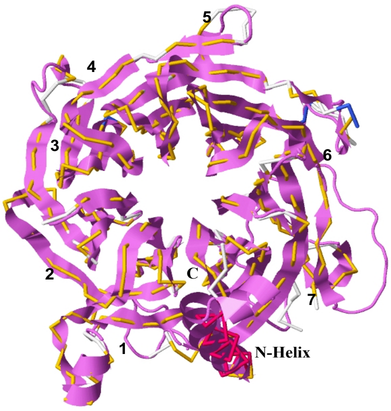


Superimposition of human RBBP4/7 and predicted structure of Tetrahymena RebL1

**Figure S5: A**: Box plot representation of RBBP4 (top) and RBBP7 (bottom) relative expression levels in normal tissues (blue) vs primary tumors (red) across multiple cancer (TCGA RNA-seq data). ** P-value cut-off ≤ 0.01(student's t-test). The TCGA cancer abbreviations are provided in supplemental Table S9. **B:** Heat map shows the survival analysis result based on multiple cancer types. The heat map depicts the hazard ratios in logarithmic scale (log10) for RBBP4 and RBBP7.The red blocks denote higher whereas blue blocks indicate lower risk, with an increase in the gene expression. The blocks with darkened frames denote statistical significance in prognostic analyses ($P \leq 0.05$; Mantel–Cox test). **C:** Histogram of gene dependency scores for RBBP4 for all cell lines in DepMap database (https://depmap.org/portal/gene/RBBP4?tab=overview) for CRISPR-Cas9 essentiality screens [CRISPR (Avana)]. Based on DepMap database, 741/757 cell lines were found to be dependent on RBBP4 expression. A lower score (CERES) means that a gene is more likely to be dependent in a given cell line. A score of 0 (indicated by black line) is equivalent to a gene that is not essential whereas a score of -1 (indicated by red line) corresponds to the median of all common essential genes. **D:** The predicted RebL1 structure shown as cartoon (violet ribbons) was superimposed with the reported human RbAp46/ RbAp48 crystal structure (PDB ID: 3CFS) depicted as gold backbone. Seven blades of predicted beta propeller are also numbered based on the human structure superimposition.

Nabeel-Shah_Figure S6

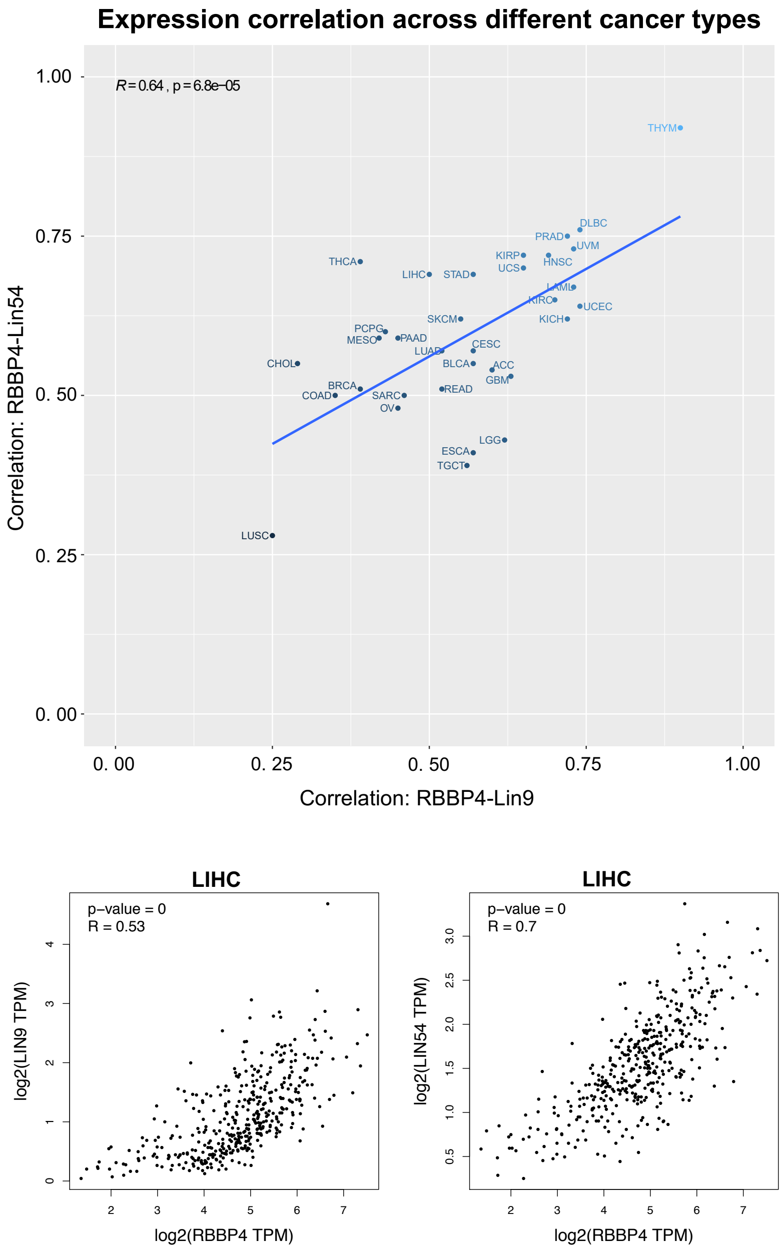

**Figure S6:** Top: Pearson correlation plot for RBBP4 to Lin54 and Lin9 expression levels across 33 TCGA cancer types. The p-value cut-off was $\leq 0.01$. The x-axis and y-axis show the correlation co-efficient values for RBBP4-to-Lin9 and RBBP4-Lin54, respectively. Each dot represents a tumor type and is labeled according to TCGA abbreviations. Overall Pearson correlation *R* is also reported along with P-value. Bottom: Representative scatter plots for RBBP4-to-Lin9 (left) and RBBP4-Lin54 (right) expression correlations. Each dot represents a single TCGA sample. 'R' denotes the Pearson correlation coefficient. TPM: Transcripts per million. The TCGA cancer abbreviations are provided in supplemental Table S9.
